# Reverse Predictivity: Going Beyond One-Way Mapping to Compare Artificial Neural Network Models and Brains

**DOI:** 10.1101/2025.08.08.669382

**Authors:** Sabine Muzellec, Kohitij Kar

## Abstract

A major goal in systems neuroscience is to build computational models that capture the primate brain’s internal representations. Standard evaluations of artificial neural networks (ANNs) emphasize *forward predictivity*—how well model features predict neural responses—without testing whether model representations are themselves recoverable from neural activity. Here we develop the *reverse predictivity* metric, which quantifies how well macaque inferior temporal (IT) cortex responses predict ANN unit activations. This two-way framework reveals a striking asymmetry: models with high forward predictivity (∼50% variance explained) often contain units unpredictable from neural activity, reflecting biologically inaccessible dimensions. In contrast, monkey-to-monkey mappings are symmetric, confirming that the asymmetry reflects genuine representational mismatch. Reverse predictivity isolates “common” ANN units—shared with IT, behaviorally relevant, and generalizing across species—and “unique” units lacking such alignment. Influenced by feature dimensionality, training objectives, and adversarial robustness, reverse predictivity offers a principled benchmark for guiding next-generation ANNs toward both high task performance and genuine biological plausibility.

## Introduction

Over the past decade, deep artificial neural networks (ANNs) trained on visual tasks have transformed our understanding of primate sensory systems^1,2^. These computational models, optimized primarily for image recognition tasks, have emerged as powerful hypothesis generation frameworks for neural processing in the primate ventral visual stream, particularly within the inferior temporal (IT) cortex^1,3^. Seminal studies have demonstrated striking similarities between ANN internal representations and neuronal activity in macaque V1^4^, V2^5^, V4^1,6,7^, IT^1,6,8^ cortices, as well as human visual^9–12^, auditory^13,14^ and language areas^15,16^, underscoring their utility in neuroscience. Indeed, state-of-the-art ANNs consistently account for roughly half of the explainable variance in IT neural responses^5^, reflecting their ability to capture essential aspects of the brain’s representational structure.

Despite these promising results, significant gaps remain in fully understanding the extent and nature of model-brain alignment. The conventional metric^1,5^ for evaluating model-brain correspondence—referred to here as forward predictivity—focuses solely on how well ANN features predict neural responses through a permissible linear transformation. Forward predictivity, therefore, is inherently a one-way comparison: it tests how well model units predict neurons, but not the converse. If an ANN were a perfect model of IT, the correspondence should be bidirectional – one ought to be able to map from model units to neurons and also from neurons to model units with comparable fidelity. However, this assumption has rarely been tested explicitly within ANNs or primates. Prior efforts^17,18^ have typically optimized models to maximize forward predictivity, implicitly treating the brain-to-model mapping as given. There has been growing recognition that two representational spaces might align well in one direction yet differ in dimensionality or information content^19–21^. In other words, ANN models and brains might not share all features, leading to an asymmetry in their relationship that forward metrics alone cannot reveal^22^.

Specifically, models may possess numerous units whose activations bear no clear correspondence with neural responses, yet still achieve comparable forward predictivity scores because a subset of other units from the same layer of the model effectively aligns with the neural data. Consequently, forward predictivity alone is insufficient for identifying models that most accurately reflect the true representational architecture of the primate brain. To probe this issue, we develop and test the concept of reverse predictivity – a metric that flips the conventional analysis. Instead of asking how well an ANN’s features can linearly predict neural responses, we ask: Can a population of IT neurons predict the activations of ANN units? This question assesses whether the model’s representational space lies within the span of the brain’s representation (as captured by the recorded neurons). If an ANN model were truly brain-like in its representations, one might expect that any unit in the model corresponds to some combination of neurons in IT. Conversely, if the model contains feature dimensions that the brain does not use, those model units would be unpredictable from neural activity – revealing a representational asymmetry.

Using large-scale neural recordings from macaque inferior temporal (IT) cortex^8^, we systematically probe representational asymmetries across a diverse suite of ANN architectures—including classic convolutional networks (e.g., AlexNet^23^), deeper feedforward CNNs (e.g. ResNet-50^24^, Inception-v3^25^), vision transformers (e.g. ViT^26^), and modern convolutional designs (e.g. ConvNeXt^27^)—with a two-way benchmark that complements forward predictivity (ANN to brain) with reverse predictivity (brain to ANN). Within this framework we distinguish common units, whose activations are reliably recoverable from IT and consistently support neuron prediction, from unique units, whose activations show little alignment with neural activity. To biologically calibrate the benchmark, we compare monkey to ANN mappings with monkey to monkey mappings, in which one animal’s IT population predicts the other’s. The latter are markedly more symmetric, confirming that the observed asymmetries are characteristic of model-to-brain, not brain-to-brain comparisons. Reverse predictivity exposes substantial mismatches that forward metrics alone do not reveal, and it varies systematically with factors such as effective dimensionality^28^, adversarial robustness^29^, and ImageNet accuracy^30^, often leading to divergent inferences from forward versus reverse evaluations. Collectively, these results motivate a bidirectional standard for model–brain assessment: reverse predictivity serves both as a diagnostic of hidden misalignment and as a design target, guiding the reduction of extraneous unique dimensions and promoting ANN representations that are more biologically plausible and behaviorally predictive.

## Results

As outlined above, complimentary to the standard practice of predicting neurons from ANNs^1,5,8^ in this study we probe how symmetric this relation is across models and monkeys. Specifically, as detailed in the sections below, we highlight the limitations of inferring ANN-brain alignment from forward predictivity and augment this metric by asking how well can monkey neurons recorded across the IT cortex predict ANN-IT units.

### Forward predictivity masks differences in model-brain alignment

The dominant approach for comparing ANNs to biological vision systems is forward predictivity—the degree to which model features can be used to predict neural responses^1,8^. This technique (schematized in **Figure 1A**) involves learning a linear mapping from model features to neural activity and has become the standard benchmark for evaluating model–brain alignment^5^. Yet as more high-performing models are developed, often optimized for classification benchmarks like ImageNet^30^, their forward predictivity scores have begun to plateau^5^ (**Figure S1**). Across architectures—from convolutional networks to transformers— percentage explained variance (%EV) in macaque inferior temporal (IT) cortex typically hovers around 50% (**Figure S1**). To illustrate an important caveat that is present within these seemingly equal high forward predictivity, we show the comparison of two representative ANNs: DenseNet-161^31^ and VGG16^32^. For both models, we extracted features from an IT-aligned layer (based on brain-score^5^, see Methods for Layer Selection), trained a linear regression model to predict the neural responses (Monkey 1: 169 sites, Monkey 2: 107 sites) from these features, and computed a cross-validated, noise-corrected percentage explained variance (%EV; see Methods for Forward Predictivity Analysis) for each recorded macaque IT neuron. Both models achieved nearly identical performance (*DenseNet-161: Median = 54%* ±*10.74 (MAD); VGG16: Median = 53%* ± *10.22 (MAD)*), with overlapping distributions of %EV (**Figures 1B–C**; *t(212) = −0.815, p = 0.416*). At first glance, these results suggest that the models are similarly brain-like. However, dissecting how they achieve their performance revealed stark differences. As shown schematically in **Figure 1A**, DenseNet-161 contains fewer units that do not contribute to the regression compared to VGG-16 (which contained a larger set of non-contributing units). This was confirmed by comparing the distributions of normalized regression weights (**Figures 1D–E**). We observed that the median normalized regression weight for VGG16 (*Median = 0.09* ±*0.05 (MAD)*) was lower compared to DenseNet161 (*Median = 0.24* ±*0.08 (MAD)*). A Wilcoxon rank sum one-tailed test confirmed the statistical difference between both distributions (*Z = −252.105, p < 0.001*). Given these observations, we hypothesized that units with consistently strong contributions across neurons reflect common representational components shared with the brain, whereas units with near-zero weights encode more unique dimensions (idiosyncratic to the ANNs). Thus, while forward predictivity captures how well a model predicts neural activity, it does not reveal which units drive that performance. Two models can have identical forward predictivity scores yet rely on fundamentally different internal strategies—differences that are biologically distinct and might be functionally meaningful.

**Figure 1.**
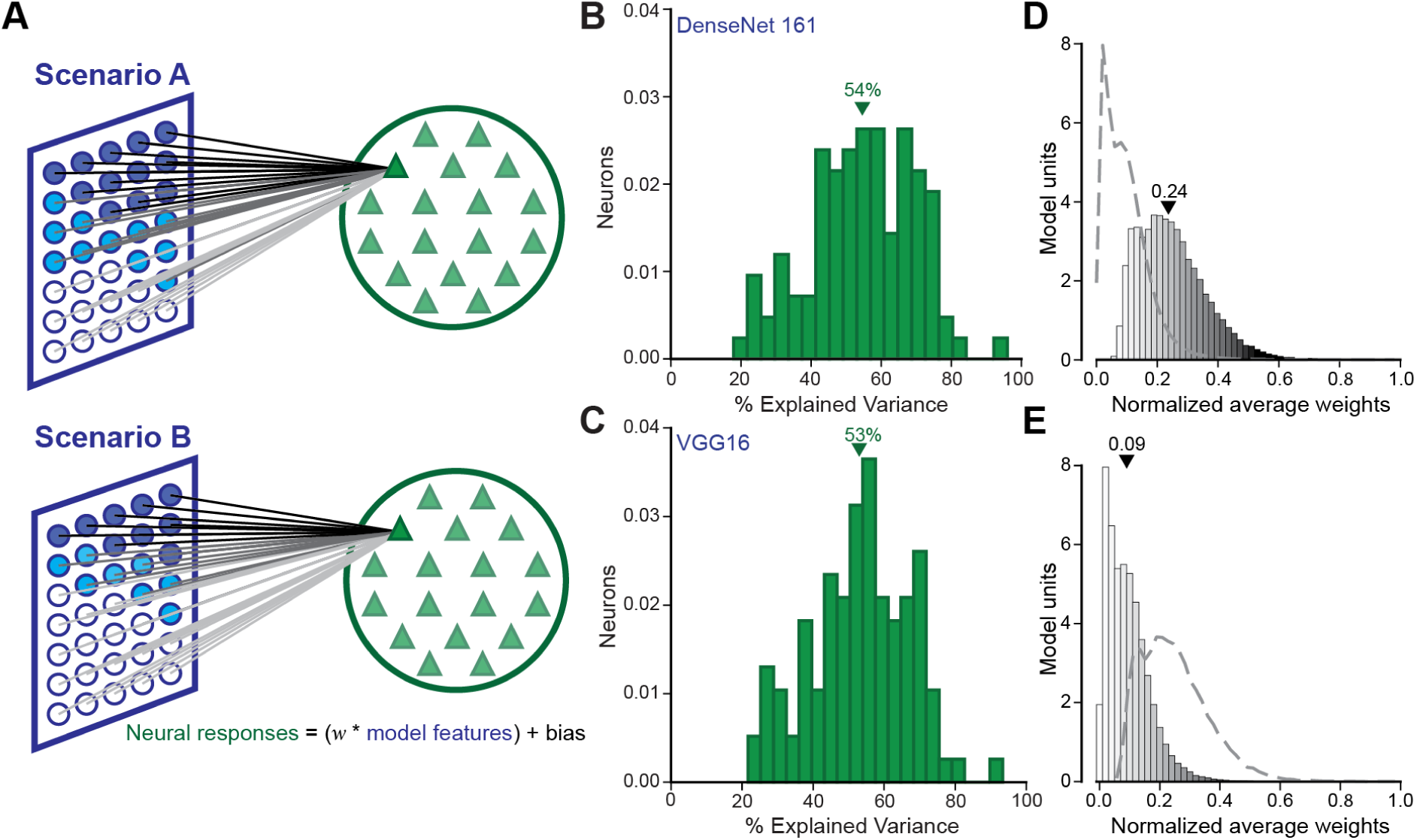
Similar predictive performance, different underlying strategies across models. (**A**) Schematic illustration of two scenarios for mapping ANN features (left) to neural responses (right), highlighting differences in unit contribution. Line darkness indicates the contribution strength of the learned linear mapping from a model unit to a neuron. **Scenario A** (e.g., DenseNet-161) shows that many units are contributing, while **Scenario B** (e.g., VGG16) exhibits more non-contributing units. (**B–C**) Distributions of explained variance (EV) across neurons when predicted by DenseNet-161 (**B**) or VGG16 (**C**) features. (**D–E**) Histograms of normalized averaged regression weights assigned to each model unit for DenseNet-161 (**D**) and VGG16 (**E**). The dashed lines indicate the weight distributions of the other model.

### Reverse predictivity reveals asymmetries between models and brains

To address the limitations of forward predictivity, we introduced a complementary metric: *reverse predictivity*. Rather than asking whether model features can predict neural activity, we asked whether neural responses can predict the activations of individual model units (**Figure 2A**). This reverses the direction of inference and probes whether a model’s internal features are linearly recoverable from brain activity. We implemented reverse predictivity by training a linear regression model for each unit in a model’s IT-aligned layer, using IT responses as input (see Methods for Reverse Predictivity). The model’s reverse predictivity score was defined as the median %EV across its units. If the differences in forward mapping weights discussed earlier genuinely reflect representational misalignment with the brain, we would expect ANNs that contain more units with higher forward weights to also be more predictable from IT responses (i.e. a higher reverse predictivity), and vice versa. To test this, we compared the average reverse predictivity scores to the average forward regression weights across ANNs (**Figure 2B**). Overall, we observed a significant positive correlation between the two measures (Spearman *r(62) = 0.4, p = 0.001,* **Figure 2B**). This relation was much stronger, when we looked at ANNs clustered according to their architectures. Specifically, we observe a strong correlation within convolutional models (Spearman *r(46) = 0.51, p<0.001*), suggesting that within well-matched architectures, reverse predictivity provides a reliable means of identifying which model features are most functionally aligned with neural representations. Consequently, reverse predictivity offers a more direct, unit-centric assessment of model–brain alignment, enabling us to classify individual units as “*common”* (high reverse %EV) or “*unique”* (low or near-zero reverse %EV). Returning to DenseNet-161 and VGG16, we observed a clear divergence in reverse predictivity. Despite similar forward scores, DenseNet-161 (**Figure 2C**) exhibited a broad distribution of reverse %EV, while VGG16’s distribution (**Figure 2D**) was heavily skewed toward zero. This difference was highly significant (Wilcoxon rank sum one-tailed test: *Z = −252.105, p < 0.001*), with average reverse predictivity score substantially higher for DenseNet-161 (*Median = 16.87, MAD = 5.96*) than for VGG16 (*Median = 5.66, MAD = 2.57*). These findings are consistent with the forward weight analyses, and confirm that similar forward predictivity scores can mask deep representational disparities between models and brains.

**Figure 2.**
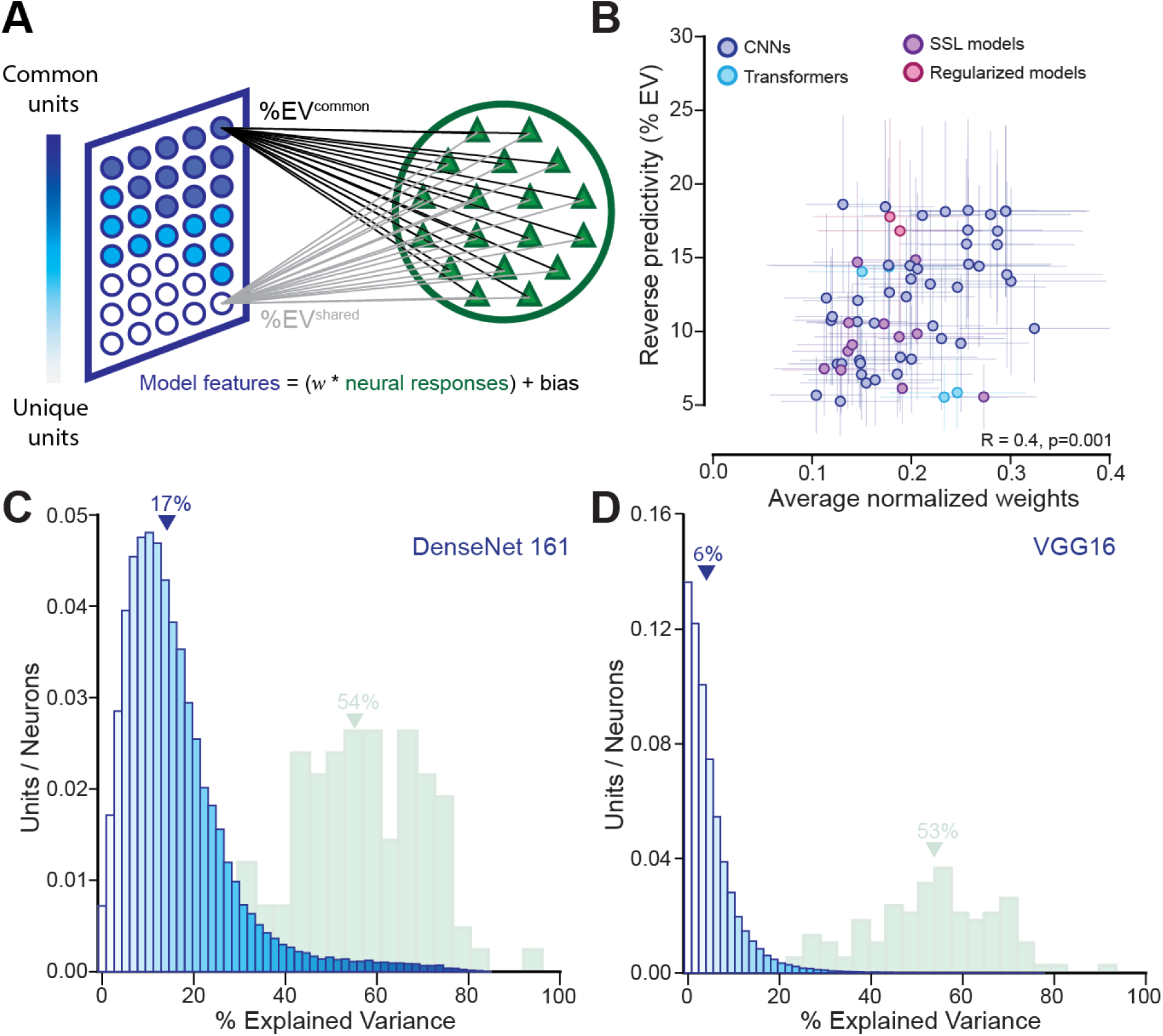
Reverse predictivity reveals differences in model-to-brain mapping strategies. (**A**) Reverse predictivity quantifies how well individual model units can be predicted from neural responses. Units with high reverse EV are considered “common”, while low reverse EV indicates “unique” units. (**B**) Reverse predictivity vs. average normalized weights across ANN models (CNNs, Transformers, SSL, robust and regularized models). Error bars indicate Median Absolute Deviation. (**C–D**) Comparison of forward (green) and reverse (blue) predictivity distributions for DenseNet-161 (**C**) and VGG16 (**D**).

We next computed forward and reverse predictivity scores for a diverse set of architectures—covering CNNs, transformers, self-supervised models, and adversarially robust variants (see Methods for Models and **Figure S4** and **Table S6** for the list of models).

We observed two striking results. First, reverse predictivity scores are significantly lower than the corresponding forward predictivity scores (Wilcoxon signed rank one-tailed test: *Z = 3240, p < 0.001*). Second, we observed a consistent inverse relation: as forward predictivity increased, reverse predictivity tended to decrease (**Figure 3**; Spearman *r(76) = –0.33, p = 0.002*;). We speculate that strategies that over the course of time have yielded higher imageNet accuracies, thereby improving forward predictivity (to a certain limit) might have introduced large mismatches between the representational structures of these ANNs with the brain – not yet accessible with forward predictivity based metrics. However, such asymmetries might be a feature of complex systems. Therefore, to first rule out this asymmetry was not common to all area-to-area comparisons across brains or machines, we applied the same analysis to monkey–monkey mappings. Interestingly, when using one monkey’s IT activity to predict the other’s, forward and reverse predictivity were closely matched (green dots in **Figure 3**), with scores along the unity line (Monkey 1 predicting Monkey 2 yielded *forward predictivity: Median = 35.5 ± 7.97 (MAD), reverse predictivity: Median = 43.2 ± 10.5*, and vice versa). This symmetry confirms that the observed model–brain asymmetry reflects a genuine representational mismatch. Furthermore, we also observe the same model-brain asymmetry and monkey-monkey symmetry when using a one-to-one mapping^33^, instead of the typical many-to-one regressions (see Methods for One-to-one mapping, **Figure S2**).

**Figure 3.**
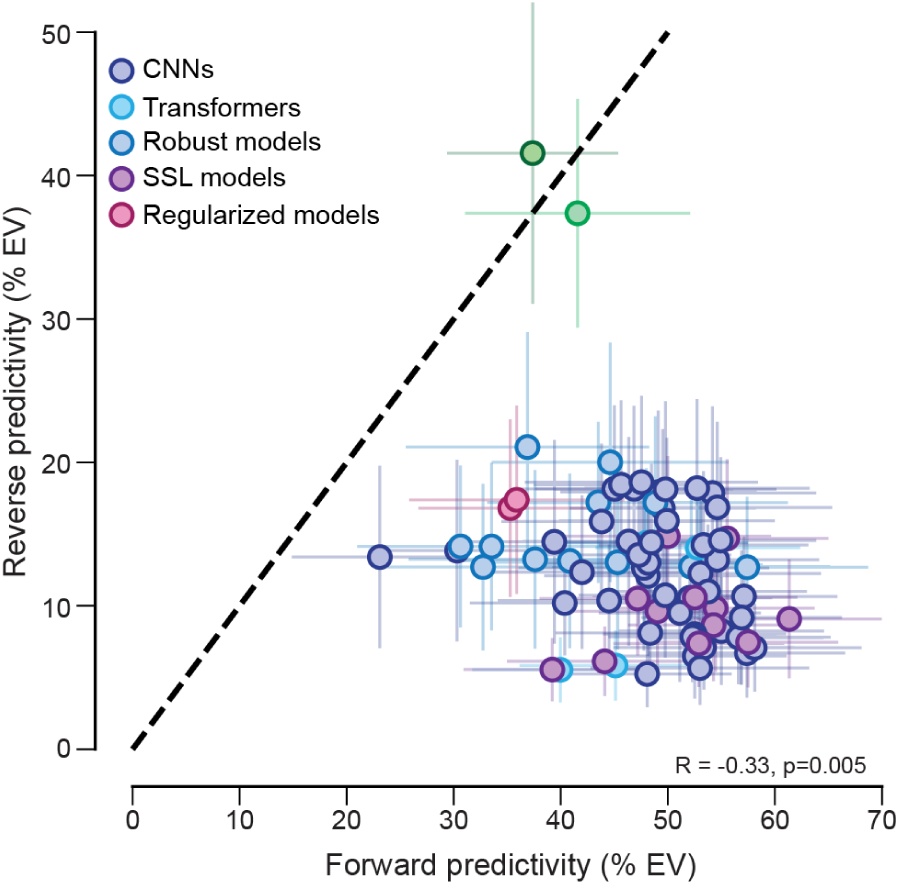
Reverse and forward predictivity do not align across models. Each dot represents a model-monkey pair, spanning a range of architectures and training regimes (CNNs, Transformers, self-supervised, robust, and regularized models). Green dots denote *Monkey-to-Monkey* predictivity, while blue to red-colored dots correspond to *ANN-to-Monkey* (forward) vs. *Monkey-to-ANN* (reverse) predictivity. Error bars represent the Median Absolute Deviation.

### Model architecture and training influence representational alignment with the brain

Next, we asked whether certain choices in model development determine a model’s reverse predictivity. Therefore, we examined how certain architectural choices, the dimensionality of the ANN-IT layers, and the training objectives affect the macaque IT’s predictivity of the ANN’s units. First, we found that while the total parameter count of the ANNs (see Methods for Model Parameter Count) had no significant effect (**Figure 4B**; *Spearman r(76) = –0.15, p = 0.237*), models with more units in their IT-aligned layer tended to have lower reverse predictivity (**Figure 4A**; *Spearman r(76) = –0.28, p = 0.012*). This suggests that feature width—not overall model size— could drive the emergence of brain-misaligned units. Second, we examined each model’s effective dimensionality^28^—quantified as the participation ratio of the PCA eigenvalue spectrum, which reflects how many dimensions are effectively used to represent visual information (see Methods for Effective Dimensionality).

**Figure 4.**
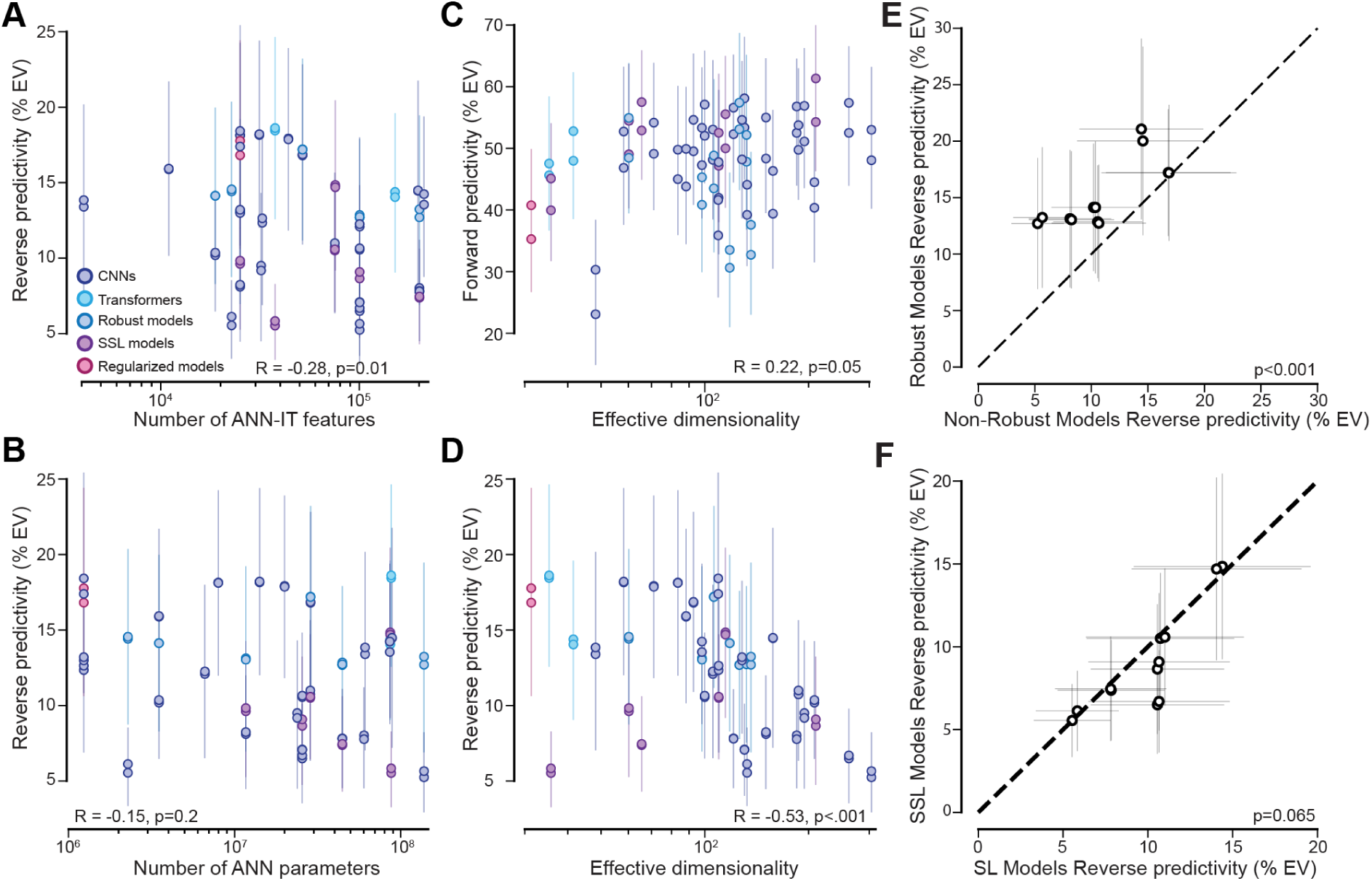
Factors influencing Reverse Predictivity. (**A–B**) Reverse predictivity across models as a function of the number of features in the IT layer used for mapping to neural data (**A**) and total number of model parameters (**B**). (**C–D**) Forward (**C**) and reverse (**D**) predictivity against effective dimensionality of model representations. (**E**) Reverse predictivity for self-supervised (SSL) vs. supervised (SL) models, matched in architecture. (**F**) Same comparison for robust vs. non-robust model pairs. Error bars represent the Median Absolute Deviation across units per model.

Consistent with previous results^28^, we indeed observed that forward predictivity increased with effective dimensionality (**Figure 4C**; *Spearman r(76) = 0.22, p = .05*). In contrast, reverse predictivity decreased with dimensionality (**Figure 4D**; *Spearman r(76) = –0.53, p < .001*), indicating that high-dimensional models accumulate features not present in IT. Third, we asked whether learning strategies that are known to reduce the dimensionality of ANN-IT features, yielding a better representational match with macaque IT neurons^34^, could improve reverse predictivity. Therefore, we tested the effects of adversarial training to generate more robust models on reverse predictivity. Indeed, adversarially robust models consistently outperformed their non-robust counterparts in reverse predictivity (**Figure 4F**; *t(11) = 5.762, p<.001*). Thus robust training appears to regularize models toward more biologically compatible features. Finally, we further explored the role of learning strategies during model training. Comparing supervised and self-supervised models with identical architectures, we observed a trend toward higher reverse predictivity in supervised models (**Figure 4E**; *t(11) = 2.07, p=0.06*). While not statistically significant, the effect suggests that SSL may induce less brain-aligned representations. In addition, while we find a positive correlation between forward predictivity and ImageNet^30^ top-1 accuracy (*Spearman r(76)=0.69, p<0.001*), we observed a trend towards a negative correlation between reverse predictivity and ImageNet top-1 accuracy (*Spearman r(76)= −0.25, p=0.12*). Together, these results show that reverse predictivity is sensitive to certain ANN design choices, not aligned with improving forward predictivity.

### Unique units reflect genuine representational mismatches—not sampling limitations

Could unique units simply reflect noisy data or limitations in neural sampling during recordings? To rule this out, we first tested whether ANN unit predictability was consistent across animals. We computed reverse %EV for each ResNet-50 unit using two monkeys separately. The results were highly correlated (*Spearman’s r(100350) = 0.87, p < .001*), suggesting a strong agreement in unit ranking (**Figure 5A**, see also **Figure S3A** for consistent correlations across all tested models). Next, we tested whether low predictivity arose from undersampling. Focusing on the 500 least predictable ResNet-50 units, we subsampled neurons from 20 to 300 and assessed reverse EV (Figure 5B left, see also *Sampling Analysis of Reverse Predictivity*). Performance plateaued quickly (with maximal reverse predictivity scores of: *Monkey 1=2.87 %EV* ± *0.07 (MAD)* and *Monkey 2=2.85 %EV* ± *0.06 (MAD)*), suggesting that even with more neural samples, these units remain unpredicted by the population of macaque IT. The responses of the unique units however were not inherently unpredictable. They were highly predictable from other model units—especially within the same model instance (**Figure 5C**, dark blue dots), but also from other instances (pre-training weight initializations) of the same model (**Figure 5C**, lighter blue dots). We observed that predictability of the unique units was lower for different random initializations of the same ANN architectures compared to the same instance (same instance %EV > different instances %EV Wilcoxon signed rank one-tailed test: *Z = 36.0, p < .001*), indicating that these unique units encode structured but model-specific representations. This result was consistent across models with units within models being better predictive of unique units than macaque IT neurons (see **Figure S3B**).

**Figure 5.**
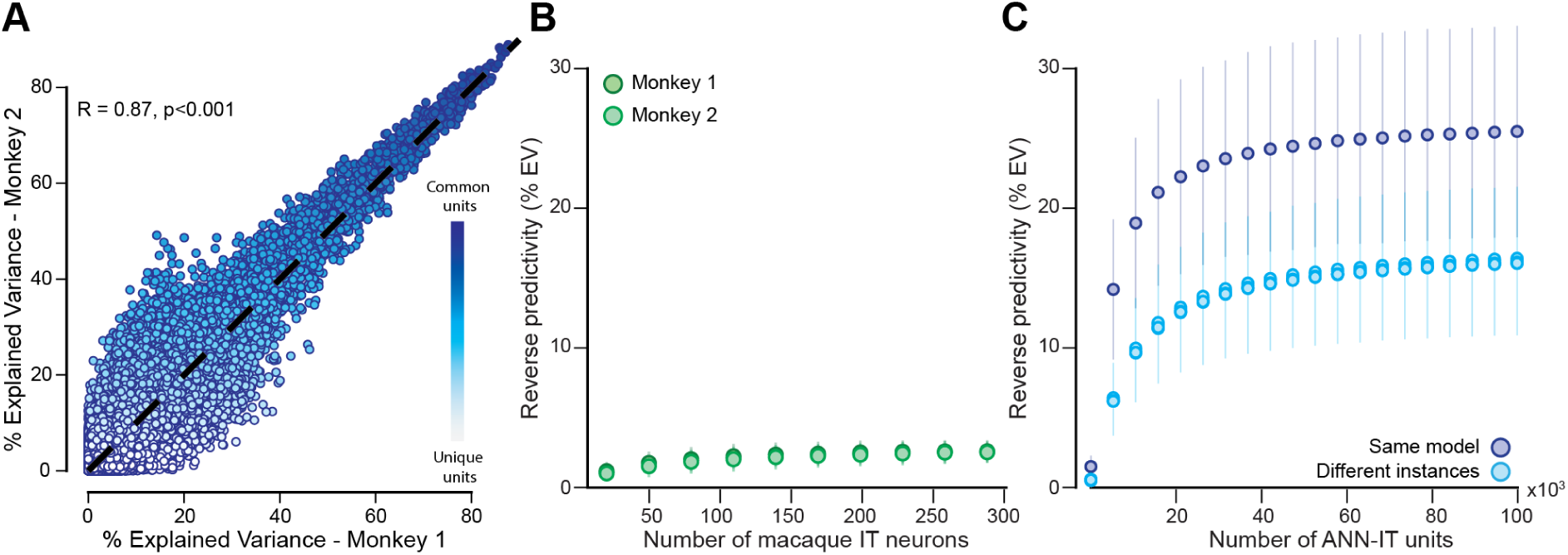
Unique units are consistent across monkeys and independent of neuron sampling. (**A**) Explained variance (%EV) of model units when predicted from each of two monkeys independently. Each dot represents a unit of ResNet50. Unique units (light) are characterized by low %EV while common units (dark blue) exhibit high %EV. **(B-C)** Sampling analysis to predict unique units. Error bars represent median absolute deviation. (**B**) Monkey-to-model reverse predictivity, as a function of sampled monkey neurons for either monkey. (**C**) Reverse predictivity given units sampled from the same layer of ResNet50l (dark blue), or the same layer of different instances of ResNet50 (light blue).

### Common units across models are better predictors of downstream behavioral variance across species

To investigate the functional significance of the common and unique units, we next computed the ability of these units to predict object discrimination behavior across various models and primates. We selected the top (*common*) and bottom (*unique*) 20% of ANN-IT units based on reverse predictivity, and applied a subsampling procedure to ensure that common and unique units reach comparable accuracies (see Methods for *Consistency of Common and Units with Behavioral responses,* **Figure S4C**). Then we trained cross-validated linear classifiers on responses from each unit type to decode object identity, generating image-by-image behavioral accuracies (**Figure 6A**, see Methods for *Behavioral Metrics*). These predictions were compared to those from other models and to empirical behavioral responses from three monkeys and a pool (n=88) of human participants (**Figure 6B**, see *Consistency of Common and Unique Units with Behavioral Responses*). Because common units are selected based on their alignment with IT cortex responses—a region that both supports core object recognition and robustly predicts behavioral performance^35^—they are expected to be more behaviorally relevant, reflecting task-relevant visual representations.

**Figure 6.**
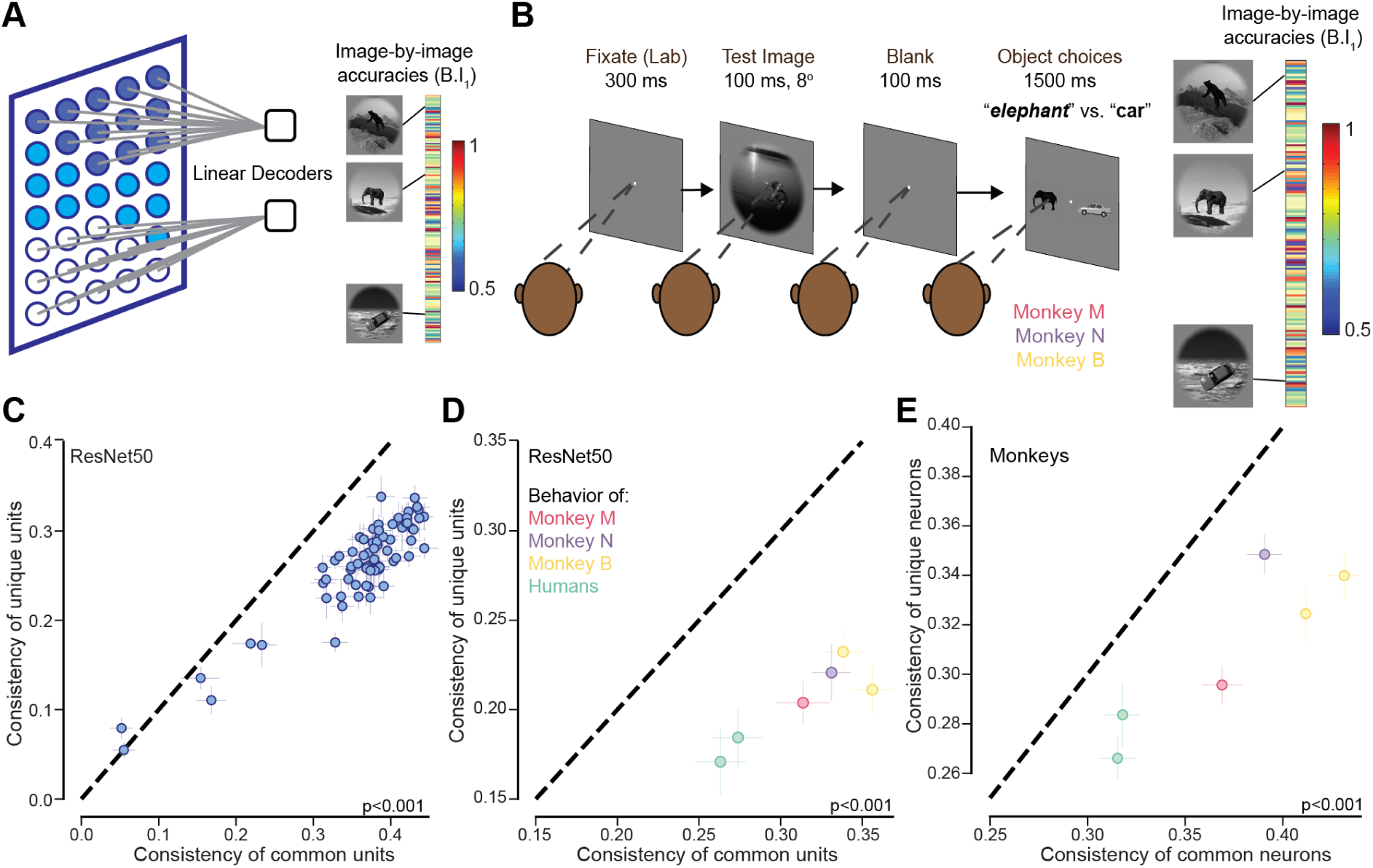
Common units drive consistent behavior prediction across models and brains. (**A**) Schematic of model-based behavioral prediction. Linear decoders are trained on common and unique units to predict object categories for each image, yielding an image-by-image behavioral signature. (**B**) Schematic of the behavioral paradigm used with monkeys: after a brief fixation, test image, and delay, animals choose between two object categories (e.g., *elephant* vs. *car*). Behavioral signatures are generated similarly for three monkeys and humans. (**C**) Model-model consistency: consistency of common units vs unique units of ResNet50 with all other models’ behavior. (**D**) Model-to-brain consistency: consistency of common units vs unique units of ResNet50 with primate behavior (three monkeys and humans). (**E**) Monkey-to-monkey consistency: consistency of common neurons vs unique neurons of two monkeys with primate behavior (same two monkeys as well as a third held-out monkey and humans). Error bars represent median absolute deviation.

In all comparisons, common units outperformed unique ones. Their behavioral predictions were more consistent with other models (**Figure 6C**; Wilcoxon signed rank test*: Z = 3081.0, p < .001*) with common units achieving higher consistency (*Median = 0.43* ± *0.009*) compared to unique units (*Median = 0.35* ± *0.004*). Interestingly, common units are also more aligned with primates’ (both monkeys and humans) behavior (**Figure 6D**; *t(7)=5.71, p<0.001*). This trend held across architectures (**Figure S4**) and also when common and unique units were selected based on one-to-one mapping (**Figure S5**). Furthermore, common ANN units identified using neural activity from two monkeys separately, better predicted behavior in other monkeys and humans than unique units (**Figure 6E***; t(7)=6.24, p<0.001*). Together, these results demonstrate that reverse predictivity not only identifies the model features most aligned with neural activity of macaques, these units are also most predictive of object discrimination behavior, across species and systems.

## Discussion

Our findings establish a fundamental asymmetry between current ANN models and the primate visual cortex. While existing evaluations have focused on forward predictivity^1,5^—how well model features can approximate neural responses—we demonstrate that this one-way comparison fails to identify truly brain-like models. The introduction of reverse predictivity, which quantifies how well IT neural activity can recover ANN unit activations, reveals that many model features remain untethered from any neural counterpart. This asymmetry was evident across model architectures and stands in contrast to the symmetry observed in monkey-to-monkey mappings, indicating that the discrepancy reflects genuine representational mismatch rather than noise or sampling limitations.

This asymmetry challenges the assumption that better forward predictivity implies a model’s internal representations are more biologically plausible. As we demonstrate (**Figure 1, 2**), two models can achieve comparable forward scores while relying on strikingly different strategies—some with many units tightly aligned to neural activity, others with large pools of dimensions that serve no evident biological purpose. Reverse predictivity exposes these hidden differences, flagging the presence of units that, although useful for the model’s task, are unaccounted for in the primate brain. The existence of such units suggests that current ANNs achieve high task performance with computational solutions that the brain may not implement, and which are not revealed by conventional benchmarks. These unique units are not simply the product of noise or poor sampling. We observed strong consistency in unit predictability across monkeys (**Figure 5A**) and found that additional neurons did not improve the reverse predictivity of the least predictable units (**Figure 5B**). Rather, these units are readily predicted by other ANN units—especially within the same architecture—indicating that they carry structured, yet model-specific, information (**Figure 5C**). The brain’s failure to predict them thus reflects not an inability to approximate model representations, but a divergence in the nature of those representations themselves. Reverse predictivity also offers a functional lens: units that are better aligned with neural activity tend to yield behavioral predictions that are more consistent across species and models. The units that are predictable from IT neurons—common units—not only generalize better across architectures but also track primate behavior more closely (**Figure 6**). This suggests that the shared subspace between ANNs and IT cortex captures the representational core necessary for object recognition^36^. In contrast, unique units—though they may contribute to model performance (**Figure S4D, S5D**)—do not appear to support biologically plausible or behaviorally relevant computations. Identifying and prioritizing these shared components could thus refine both our understanding of biological vision and the development of more brain-like models.

The implications of this two-way framework extend beyond benchmarking. Reverse predictivity provides a diagnostic tool for evaluating whether model representations lie within the span of the brain’s representational geometry. Models that include extraneous dimensions—those orthogonal to IT neural activity—are inherently misaligned, even if they perform well on behavioral tasks or forward mappings. Forward and reverse predictivity together reveal a tension: increases in one metric do not guarantee increases in the other. In fact, we observe that models with higher forward scores often fare worse in reverse predictivity, pointing to a growing divergence between ANN representations and the neural code as models scale in complexity and accuracy – something that the field has to contemplate further^37^.

Factors such as feature dimensionality, training objective, and architectural design all influence reverse predictivity (**Figure 4**). Larger representational spaces tend to dilute neural alignment, while adversarial training improves it. Interestingly, self-supervised learning—despite its recent success—tends to yield lower reverse predictivity than supervised training. These insights suggest that reverse predictivity can serve not only as a benchmark but also as a constraint for future model development. Penalizing dimensions not recoverable from neural activity may help enforce tighter alignment between artificial and biological systems. In addition, rather than optimizing only for task performance or forward neural predictivity, future models could be explicitly optimized to maximize ensemble common unit match—training ANNs such that their internal units become recoverable from other ANN units. Such alignment-based training may help reduce idiosyncratic units and yield more brain-compatible models without sacrificing performance.

Reverse predictivity also complements prior approaches such as representational similarity analysis^40,41^. While those methods capture global representational geometry or shared subspaces, reverse predictivity directly quantifies whether the full span of model features is embedded within the brain’s representational structure. The asymmetries we uncover suggest a many-to-one mapping: the brain captures only a subspace of model features, and the inverse mapping cannot recover the full model representation. This implies a non-invertible relationship—one in which ANN representations are richer or at least distinct from those in the IT cortex.

While our analyses focused on the IT cortex, it remains possible that some unique units correspond to computations in other brain regions^42^. However, the symmetry observed in monkey-to-monkey mappings, and the consistent behavioral alignment of common units, argues that the primary dimensions used for core object recognition are well captured by our IT recordings. Increasing the number of neurons or expanding the stimulus space might uncover additional alignments, but our sampling analyses suggest diminishing returns. Likewise, the assumption of linear mappings could be revisited, but if model features require non-linear combinations of neural responses to be recovered, that itself implies a mismatch in representational format.

Similarly, although our study was grounded in vision, the conceptual framework is broadly applicable. In neuroscience domains where ANN–brain comparisons are now commonplace—such as audition^13^, language^15^, and motor control^38^—reverse predictivity provides a principled tool to test whether model representations are truly embedded in the corresponding neural codes. Just as forward metrics have helped map task-driven ANNs onto sensory and motor areas, reverse predictivity can help ensure that such mappings are not only functionally relevant but structurally faithful. This is especially relevant as ANN-based models are increasingly used to interpret neural data and propose mechanistic hypotheses across cortical systems^39^.

Ultimately, the goal of neuroscience-aligned modeling is not merely to match neural outputs, but to mirror the brain’s internal structure and computations^3^. Reverse predictivity advances this goal by assessing whether a model’s internal code is embedded in neural activity. Our findings demonstrate that no current ANN fully meets this criterion—each carries representational baggage that the brain does not use. A model with high forward and reverse predictivity would reflect both behavioral performance and biological plausibility. We anticipate that reverse predictivity will become a critical target in the development and selection of next-generation models, guiding architectures toward representations that not only solve tasks but also resonate with the brain’s intrinsic structure. To facilitate systematic model comparison, we propose a unified metric—the *Bidirectional Predictivity Index (BPI)*—defined as the harmonic mean of forward and reverse predictivity. This metric rewards models that not only predict neural responses well but are also themselves predictable from neural activity. Specifically, given a model’s average forward predictivity score (F) and reverse predictivity score (R), the index is computed as: *BPI*=2*F*R/(F+R). This formulation penalizes asymmetries and provides a single interpretable score for ranking models by their overall representational alignment with the brain. Unlike forward predictivity alone, which can mask large sets of brain-inaccessible units, or reverse predictivity alone, which may undervalue functionally useful representations, the BPI captures the extent to which a model’s internal space is both sufficient and necessary for capturing brain responses. Future work could extend this framework by normalizing BPI relative to brain-to-brain ceilings or incorporating penalties for dimensionality mismatches. As ANN–brain comparisons grow across sensory and cognitive domains, such a unified score can serve as a principled benchmark for building models that truly reflect biological computation.

## Methods

### Visual Stimuli

We used 1320 images previously used in a study^8^. The images contained 10 objects types (bear, elephant, face, apple, car, dog, chair, plane, bird, zebra). Images were either synthetic (but naturalistic) or natural photographs from the MS COCO imageset^43^. The synthetic images were rendered from a 3D object model (originally purchased from TurboSquid) into a 2D projection while varying an object’s position (horizontal and vertical position), rotation (in the horizontal, vertical, or depth plane), and size. The rendered object view was then added to a randomly chosen natural image background (indoor and outdoor scenes obtained from Dosch Design, www.doschdesign.com). All resulting images were gray-scaled and had a resolution of 256×256 pixels. The MS COCO images were colored but also contained the objects from the 10 categories mentioned above. We used a high-resolution display with a refresh rate of 60 Hz for neural recordings and standardized luminance across images to control for low-level visual confounds.

### Subjects

#### Non-human primates

The nonhuman subjects in our experiments were three adult male rhesus monkeys (Macaca mulatta). All surgical and animal procedures were performed in accordance with National Institutes of Health guidelines, the Massachusetts Institute of Technology Committee on Animal Care, and the Canadian Council on Animal Care on the use of laboratory animals and were also approved by the York University Animal Care Committee.

### Large-Scale Neural Recordings in the Inferior Temporal Cortex

Data analyzed in this article has been previously used for various scientific publications^8,44,45^.

#### Surgical Implants and Microelectrode Arrays

We surgically implanted three 10×10 microelectrode arrays (Utah arrays, Blackrock Microsystems) per hemisphere in the inferior temporal (IT) cortex of each monkey under aseptic conditions. Each array contained 96 electrodes, excluding corner electrodes, with a length of 1.5 mm and a spacing of 400 μm between electrodes. We determined the placement of the arrays intraoperatively using the visible sulcus patterns for guidance. For monkeys receiving implants in both hemispheres, we initially implanted arrays in one hemisphere and recorded data for approximately one year before explanting and reimplanting new arrays in the opposite hemisphere.

#### Electrophysiological Recordings

During each experimental session, multiunit neural activity was recorded continuously at a sampling rate of 20 kHz using an Intan RHD Recording Controller (Intan Technologies, LLC). The raw voltage signals were bandpass filtered offline using a second-order elliptical filter (300 Hz to 6 kHz, 0.1 dB passband ripple, 50 dB stopband attenuation), before being thresholded to obtain the multiunit spike events. A multiunit spike event was defined as the threshold crossing when voltage (falling edge) deviated by more than three times the standard deviation of the raw voltage values.

The implanted arrays sampled a range of regions along the posterior-to-anterior axis of the IT cortex. For all analyses, we treated each recording site as a random sample from the broader IT population without considering the precise spatial locations of the electrodes.

#### Eye Tracking and Calibration

During recording sessions, we monitored eye movements using video eye tracking (SR Research EyeLink 1000). Using operant conditioning and water reward, our subjects were trained to fixate a central white dot (0.2°) within a square fixation window that ranged from ±2°. At the start of each behavioral session, monkeys performed an eye-tracking calibration task by making a saccade to a range of spatial targets and maintaining fixation for 500 ms. Calibration was repeated if drift was noticed over the course of the session.

Real-time eye-tracking was employed to ensure that eye jitter did not exceed ±2°, otherwise the trial was aborted, and data discarded. Stimulus display and reward control were managed using the MWorks Software (https://mworks.github.io).

### Behavioral Testing

#### Non-human Primates

In the behavioral experiment, monkeys performed an object recognition task. Images were presented on a 24-inch LCD monitor (1920 × 1080 at 60 Hz) positioned 42.5 cm in front of the animal. Monkeys were head-fixed. Monkeys fixated on a white dot (0.2°) for 300 ms to initiate a trial. The trial started with the presentation of a sample image (from a set of 1320 images) for 100 ms at approximately 8 degrees of visual angle. This was followed by a blank gray screen for 100 ms, after which the choice screen was shown, containing a standard image of the target object (the correct choice) and a standard image of the distractor object. The monkey was allowed to view freely the choice objects for up to 1500 ms and indicated its final choice by holding fixation over the selected object for 400 ms. Trials were aborted if gaze was not held within ±2° of the central fixation dot at any point until the choice screen was shown. Prior to testing in the laboratory, monkeys were trained in their home-cages (similar to Ramezenpour et al., 2024^44^) to perform the delayed match to sample tasks on the same object categories (but with a different set of images, see Sörensen et al. 2024^45^ for details).

Behavioral performance was quantified on an image-by-image basis, as the proportion of correct responses for each image across all distractors (see *Behavioral Metrics* below for more details).

#### Humans tested via Amazon MTurk

We measured human behavior (88 subjects) using the online Amazon MTurk platform, which enables efficient collection of large-scale psychophysical data from crowd-sourced human intelligence tasks. The reliability of the online MTurk platform has previously been validated by comparing results obtained from online and in-lab psychophysical experiments^35,46^. Each trial started with a 100-ms presentation of the sample image (1 out of 1,320 images). This was followed by a blank gray screen for 100 ms followed by a choice screen with the target and distractor objects. The subjects indicated their choice by touching the screen or clicking the mouse over the target object. Each subject saw an image only once. We collected the data such that there were 80 unique subject responses per image with varied distractor objects.

#### Behavioral Metrics

We have used a one-vs-all image level behavioral performance metric, **B.I_1_**, similar to previous studies^8,46,47^, to quantify the behavioral performance of monkeys and humans. This metric estimates the overall object discriminability of each image containing a specific target object from all other objects (pooling across all 9 possible distractor choices).

As mentioned above, for each trial of the task, a specific sample image was shown, and a binary choice task screen was presented. So, the data obtained from each trial is a correct (1) or incorrect (0) choice from the subjects. Each trial can be labeled with 2 unique identifiers – the unique sample image (one out of 1320) and a unique task (one out of nine possible tasks given the image, e.g. bear vs. dog).

Hence, given an image of object ‘i’, and all nine distractor objects (j≠i) we computed the average performance per image (each element of the **B.I_1_** vector) as the average of the percent correct across all the binary tasks done with that image as the sample image (where object ‘i’ was the target and all objects j≠i were the distractors respectively).

While the **B.I_1_**vector provides an estimate of the monkeys’ image-by-image accuracy, the monkeys’ overall performance accuracy can be determined by taking an average of the **B.I_1_** vector (across all images).

#### Trial split-half reliability estimation

To compute the reliability of the B.I_1_ vector, we split the trials per image into two equal halves by resampling without substitution. The median of the correlation of the two corresponding vectors (one from each split half), across 100 repetitions of the resampling was then used as the uncorrected reliability score (i.e., internal consistency), r. To correct for the usage of half the number of trials to estimate the reliabilities in comparison to the raw correlations, we used the Spearman Brown correction method^48^ on the uncorrected reliability score (r) as follows,

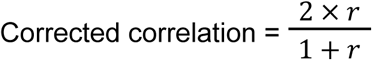

#### Noise ceiling estimation

The noise ceiling is computed as the square root of the product of the trial split half reliabilities of each of the variables used in the raw correlation. For instance, to estimate the noise ceiling (maximum correlation expected) for the comparison of monkey 1 vs. human pool B.I_1_, the two relevant internal reliabilities are those estimated for B.I_1_ of the monkey 1 and human pool respectively.

### Analyses of ANN Models

#### Model selection

We selected a broad and diverse set of 39 artificial neural network (ANN) models spanning a wide range of architectures, training objectives, and inductive biases. These models include classic convolutional networks such as AlexNet^23^, VGG16^49^, and multiple ResNet^24^ variants (e.g., ResNet-18, ResNet-50, ResNet-101, ResNet-152), as well as deeper architectures like ConvNeXt^27^, DenseNets^31^, Inception-v1/v3^50,25^, and NASNet/PNASNet^51,52^. We also included lightweight models such as MobileNet^53^, ShuffleNet^54^, and SqueezeNet^55^, biologically inspired models like CORnet-S and CORnet-RT^56^, and modern transformer-based architectures including Vision Transformers^26^ (ViT) and Swin Transformers^57^. To capture the effects of training, we incorporated models trained under standard supervised learning, self-supervised learning (e.g., SimCLR^58^, SwAV^59^), and robust models developed by adversarial training^60^. All models were pre-trained on the ImageNet^30^ object classification dataset and task (except for minor differences in training specifics as per the original publications), and we used the publicly available pre-trained weights.

#### Layer selection

For each ANN, we extracted activations from a single high-level layer corresponding to the macaque IT cortex, as defined by the ANN-IT mapping on Brain-Score^5^. The layer selection in Brain-Score is done by asking which layer of the model best predicts existing macaque IT data, using data from a previous publication^35^, and methods previously published^1^ (similar to our many-to-one regression method). Of note, we did not use the same neural data used in layer selection for our analyses in the article. Neural data^8^ used in this article was collected in a separate set of monkeys using a different image-set. This layer produces an M-dimensional feature vector per image, where M varies by model (e.g., 4096 for AlexNet, 100372 for ResNet-50). For a stimulus set of N images, this yielded an M × N matrix of model responses per ANN (See **Table S6** for a full list of models and their IT layers).

#### Effective Dimensionality

To estimate the representational complexity of each model’s IT-layer features, we computed effective dimensionality using the participation ratio of the PCA eigenvalue spectrum. For each model, given the activation matrix of units in the IT-mapped layer, we performed principal component analysis (PCA) to obtain the eigenvalues of the covariance matrix. The effective dimensionality was then defined as the participation ratio:

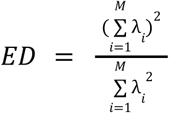

Where λ*_i_* denotes the variance explained by the *i*-th principal component. This measure reflects the number of dimensions effectively used to represent the stimulus set, taking into account the distribution of variance across components.

#### Model Parameter Count

To estimate model capacity, we computed the total number of trainable parameters for each architecture. For models implemented in PyTorch or available via the timm library, we loaded the model in an untrained state (pretrained=False) and used model.parameters() to count all trainable parameters. The total parameter count was computed as the sum of all elements in the model’s trainable tensors. For models not directly supported in PyTorch (e.g., CORnet variants), we used fixed parameter counts based on documentation or prior reports. These values were used to examine the relationship between model size and reverse predictivity (see Figure **4B**).

#### Matched Model Comparisons

To isolate the effect of the training objective, we selected model pairs with identical architectures but differing in training strategy. For self-supervised vs. supervised comparisons (**Figure 4E**), we matched 6 models such as ResNet-50 trained with SimCLR^58^ vs. supervised ImageNet^30^. For robustness comparisons (**Figure 4F**), we used architectures trained with adversarial robustness objectives^61^ and their standard counterparts. Reverse predictivity scores were averaged across units for each model and compared pairwise across models.

#### Self-Supervised vs. Supervised Comparisons

To evaluate the effect of training objective on reverse predictivity, we compared models with identical architectures but trained under supervised (SL) and self-supervised (SSL) regimes including approaches such as SimCLR^58^, CLIP^62^, or DINO^63^ (**Figure 4E**). For ResNet-50, we included two SSL variants: one trained with semi-supervised learning (SWSL^59^) and one trained with fully self-supervised objectives. These were compared against the standard supervised version. Additional matched comparisons include: ConvNeXt^27^, Vision Transformer^26^, ResNet-101^24^, and Swin Transformer^57^. For each pair, we computed the mean reverse predictivity score across model units, using the same IT-mapped layer. Comparisons were visualized by plotting SSL vs. SL scores for each architecture, with error bars reflecting median absolute deviation.

#### Robust vs. Non-Robust Comparisons

To investigate the impact of adversarial robustness on reverse predictivity, we compared standard models with their robustly trained counterparts across matched architectures (**Figure 4F**). Specifically, we included the following model pairs: DenseNet-161^31^, MobileNet-v2^64^, VGG-16^49^, ResNet-50^24^, ShuffleNet^54^, and ResNet-18^24^. All robust models were taken from the Madry Lab’s Robust ImageNet Models repository on Hugging Face (https://huggingface.co/madrylab/robust-imagenet-models)^61^ and trained with *l*2-constrained adversarial objectives using Projected Gradient Descent^29^ (PGD) with a perturbation bound of ɛ = 3. For each pair, we computed the mean reverse predictivity across model units in the IT-mapped layer and compared performance between robust and standard variants. Scatter plots reflect unit-wise averages per model, with error bars representing median absolute deviation.

### Analyses

#### Forward Predictivity Analysis

To quantify forward predictivity, we evaluated how well each model’s features could linearly predict IT neural responses, following standard procedures. We only used the most reliable neural sites (split half reliability > 0.7). Then, for each model and each recorded neuron *j*, we fit a linear regression (ridge regression) to predict the neuron’s responses across images from the model’s activations, with a cross-validation procedure (see Cross-Validation section). Given a response vector *r_j_* ∈^1×*N*^ and a model activation matrix *A* ∈*^M^*^×*N*^, we estimated weights *w*_*j*_ ∈^M × 1^ and bias *b* to minimize the squared error:

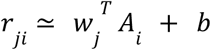

where *A_i_* is the model feature vector for a given image *i*.

Model performance was assessed using explained variance (EV), defined as the squared Pearson correlation between predicted and actual firing rates, normalized by the geometric mean of their split-half reliabilities (see Split-Half Reliability section), and scaled to percentage. We computed %EV for each neuron, of each monkey and for each model. A model’s overall forward predictivity score was defined as the mean %EV across the full population of neurons.

#### Weight Analysis

The linear weight vectors *w_j_* obtained from the regression models indicate the contribution of each model unit to the prediction of individual IT neurons. To enable comparisons across models, we normalized (division by the maximum value) the weight magnitudes within each model. We then assessed the distribution of weights for each unit across all neurons to identify units that were consistently influential versus those that were effectively ignored.

#### Reverse Predictivity Analysis

Reverse predictivity measures how well the recorded IT neuron population can predict the activations of individual units within each model’s IT-layer representation. For each model unit *k*, we treated its responses across images as the target variable and used the corresponding neural responses as predictors. Specifically, we fit a linear model of the form:

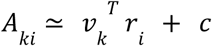

Where *A_ki_* is the activation of a model unit *k* for image *i*, *r_i_* is the vector of IT neural responses for that image, *v_k_* is a weight vector, and *c* is a bias term. We used linear regression (ridge regression) with cross-validation (see Cross-Validation section) to fit the model for each unit. Predictive performance was quantified using explained variance (%EV), computed as the squared Pearson correlation between predicted and actual activations, normalized by the geometric mean of the split-half reliabilities of the prediction and the target (see Split-Half Reliability section), and expressed as a percentage.

We computed EV for each model unit predicted from each monkey IT cortex. A model’s overall reverse predictivity score was defined as the mean %EV across all units. To characterize units by their reverse predictivity, we labeled those in the top 20th percentile as *common units* and those in the bottom 20th percentile as *unique units*.

#### Sampling Analysis of Reverse Predictivity

To examine how the number of sampled features affects the ability to predict unique units, we performed two complementary analyses. On the neural side (**Figure 5B**), we randomly subsampled increasing numbers of IT neurons from each monkey (20 to 300, in steps of 20) and evaluated reverse predictivity for the 500 least predictable units (i.e., lowest %EV) in the ResNet-50 IT layer. Each sample size was repeated 10 times, and the mean explained variance (%EV) across the 500 units was reported. On the model side (**Figure 5C**), we sampled increasing numbers of units (1,000 to 100,000 in steps of 5,000) from the same ResNet-50 layer—excluding the 500 target units to be predicted—and used them to predict these same 500 units via linear regression. We tested two conditions: units sampled from the same model instance (dark blue) and units sampled across 5 independently initialized ResNet-50 instances (light blue). In all cases, error bars represent the median absolute deviation across repetitions.

#### Monkey-to-Monkey Predictivity

To estimate an upper bound on model–brain predictivity, we evaluated how well one monkey’s IT population could predict the other’s neural responses using the same linear regression framework used for model-based analyses. For each pair of monkeys, we treated one monkey’s IT responses as predictors and the other’s as targets, using a linear regression (ridge regression) model cross-validated (see Cross-Validation section) trained across images. Predictivity was quantified as explained variance (%EV), computed per target neuron as the squared Pearson correlation between predicted and actual responses, normalized by the geometric mean of the split-half reliabilities of both the predictor and target populations (see Split-Half Reliability section). This analysis was conducted symmetrically in both directions (Monkey 1 → Monkey 2 and Monkey 2 → Monkey 1), and the resulting scores were averaged. Error bars reflect the median absolute deviation across target neurons.

#### Consistency of Common and Unique Units with Behavioral Responses

To assess how well the activity of common and unique units in models and monkeys aligned with behavioral responses, we first identified these units based on their reverse predictivity scores. Common units were selected as those in the top 20th percentile, whereas unique units comprised the bottom 20th percentile, considering predictions from both monkeys (or from the other monkey in monkey-to-monkey comparisons). To ensure comparability across models and to control for the number of units included in the behavioral analysis, we applied a random projection to reduce each unit set to 1,000 dimensions.

To control for accuracy as a potential confound, we subsampled the common units to match the accuracy level of the unique units since these units typically yield lower accuracy. Specifically, we incrementally increased the subsample size of common units (from 150 units to 350, against 1000 unique units), tracked the resulting accuracy, and selected the sample size that minimized the accuracy gap between the two groups. If the mean accuracy of the unique units fell between two subsampling values, we chose the lower-performing one for comparison. Multiple subsamples were generated following this procedure. For subsequent analyses, all unique units and subsampled common units were considered.

Behavioral responses from three monkeys and human participants were collected and summarized into image-level response vectors (see Behavioral Testing and Behavioral Metrics sections above). For each ANN model, we similarly obtained behavioral predictions by training a cross-validated linear decoder on the model’s IT-layer activations to classify image object categories (see Layer Selection and Cross-Validation sections). This procedure yielded comparable image-level response vectors for both primates and ANN models.

We repeated this decoding separately for subsets consisting exclusively of (subsampled) common or unique units, producing two additional image-level response vectors per model or monkey. The consistency between these vectors and the behavioral responses from all other models, monkeys, and human subjects was quantified using Pearson correlation. Correlations involving primate behavioral data were corrected for measurement noise by accounting for the behavioral split-half reliability (see Split-Half Reliability of Behavioral Data section below). Similarly, correlations involving neural data were corrected for neural recording noise using split-half reliability estimates specifically computed for common and unique neuron subsets (see Neural Split-Half Reliability Estimation for Common and Unique Neurons section below).

#### Neural Split-Half Reliability Estimation for Common and Unique Neurons

To estimate split-half reliability separately for the subsets of common and unique neurons, we performed the following analysis. We first partitioned neural response data (neurons × images × repetitions) into two equal subsets along the repetition dimension, independently for common and unique neuron populations. Each split-half response set was averaged across repetitions, yielding mean responses per neuron and image. Next, for each split half, we created image-level neural response vectors (**B.I_1_** vectors, see Behavioral Metrics section above). We then correlated these image-level vectors between the two splits using Pearson’s correlation coefficient. To correct for reduced reliability from splitting data in half, we applied the Spearman-Brown^48^ correction to the resulting correlation values. This procedure was repeated 10 times with randomly assigned splits to ensure robustness, and the mean correlation across repetitions was taken as the final estimate of split-half reliability for each neuron population (common and unique).

#### Split-Half Reliability of Behavioral Data

To assess the internal consistency of our behavioral measures, we computed the split-half reliability of the image-level behavioral performance metric (**B.I_1_**). For each image, we identified the minimum number of available trials across images to ensure balanced sampling. We then randomly divided these trials equally into two independent subsets (split halves) for each image. For each split half, we calculated the mean behavioral accuracy per image, resulting in two accuracy vectors (one per split half). We then computed the Pearson correlation between these two accuracy vectors across images, considering only images with valid data in both splits. This correlation coefficient served as a measure of internal consistency (split-half reliability). To obtain robust reliability estimates, this procedure was repeated multiple times with different random splits, and the median correlation value was taken as the final reliability measure.

#### One-to-One Mapping for Forward and Reverse Predictivity

To evaluate one-to-one model–neuron correspondence (**Figure S2**), we used a cross-validated correlation-based mapping strategy (see Cross-Validation). For each neuron, we identified the single best-predictive model unit using training data and measured its ability to predict test responses. Specifically, we computed the average neural response across trials and used 10-fold cross-validation to partition the data. Within each fold, we z-scored the model features and neural responses based on the training set. For each unit in the model’s IT-aligned layer, we computed its Pearson correlation with the training responses of the target neuron. The unit with the highest correlation was selected as the best-predictive unit. We then measured the correlation between this unit’s test responses and the test responses of the neuron. Explained variance (EV) was computed as the squared test-set correlation, normalized by the split-half reliability of the neuron (see Split-half reliability). This produced a %EV score for each fold, which was averaged across splits and reported as the neuron’s one-to-one forward predictivity.

For reverse predictivity, the procedure was inverted: for each model unit, we identified the best-predictive neuron using training data, then assessed how well that neuron could predict the unit’s test responses. All other steps remained identical, including normalization, correlation-based selection, and reliability correction (using the neuron’s internal reliability). This one-to-one analysis provides a fine-grained, unit-level measure of alignment between model features and neural activity.

### Statistical Analyses

All statistical analyses were performed in Python using scipy.stats. Relationships between continuous variables (e.g., reverse predictivity vs. model complexity or effective dimensionality) were assessed using Spearman correlation coefficients unless otherwise noted.

#### Statistical Testing

All statistical comparisons were guided by a consistent decision procedure based on the distributional properties and pairing of the data. First, we assessed whether each distribution was approximately normal using the Shapiro-Wilk test^65^. If both groups satisfied the normality assumption, we applied parametric tests. For paired data (e.g., within-subject or within-model comparisons), we used a paired t-test (two-tailed; degrees of freedom = n – 1). For unpaired comparisons (e.g., between models), we used an independent samples t-test (degrees of freedom = n₁ + n₂ – 2).

If at least one distribution violated the normality assumption, we used non-parametric alternatives. Specifically, we applied the Wilcoxon signed-rank test for paired comparisons and the Wilcoxon rank-sum test (equivalent to the Mann–Whitney U test) for unpaired data. All tests were two-tailed unless stated otherwise. We report exact p-values and test statistics throughout.

#### Cross-Validation

A ten-fold cross-validation scheme was used, dividing the dataset into training and testing subsets. The dataset comprised 1320 images distributed equally across ten object categories (bear, elephant, person, car, dog, apple, chair, plane, bird and zebra). This design ensured a balanced representation of each category during training and testing.

#### Split-half reliability

Split-half reliability was estimated by dividing neural responses along the trial dimension into two independent halves. For monkey-to-monkey predictions, we corrected both the ground truth and the predictions—by training separate regression models. When models were used as predictors or predicted targets, we treated them as deterministic and assigned a noise ceiling of 1, without applying any correction. Reliability for each neuron was quantified using the Pearson correlation between the two splits, with final values adjusted using the Spearman-Brown^48^ formula.

## Acknowledgments

KK has been supported by funds from the Canada Research Chair Program, the Simons Foundation Autism Research Initiative (SFARI, 967073), Brain-Canada Foundation, the Canada First Research Excellence Funds (VISTA Program), and the National Sciences and Engineering Research Council of Canada (NSERC). SM is funded by Connected Minds Postdoctoral Fellowship (supported by CFREF).

## Supplementary Figures

**Figure S1.**
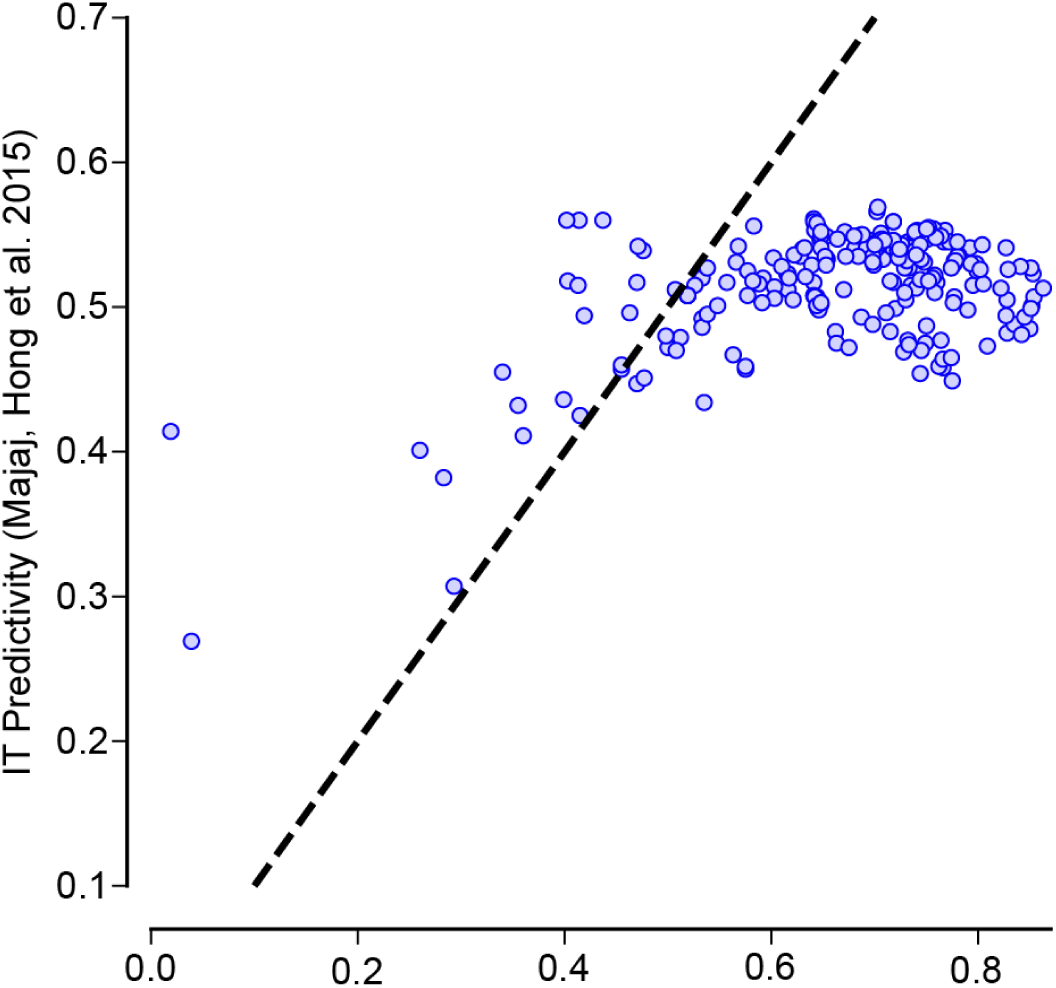
Evolution of ImageNet performance vs. forward predictivity. Scatter plot showing the relation between ImageNet top-1 classification accuracy and forward predictivity for 254 ANN models in the MajajHong2015 IT benchmark^35^. Each dot represents a model. Spearman correlation coefficient (*Spearman r(254) = 0.15, p<0.024*) indicates a moderate relationship between task performance and alignment with IT responses. However, as models get more accurate on ImageNet, their forward predictivity plateaus around 0.5 (i.e. approx 50 % EV). Data for the plot is adapted from http://brain-score.org.

**Figure S2.**
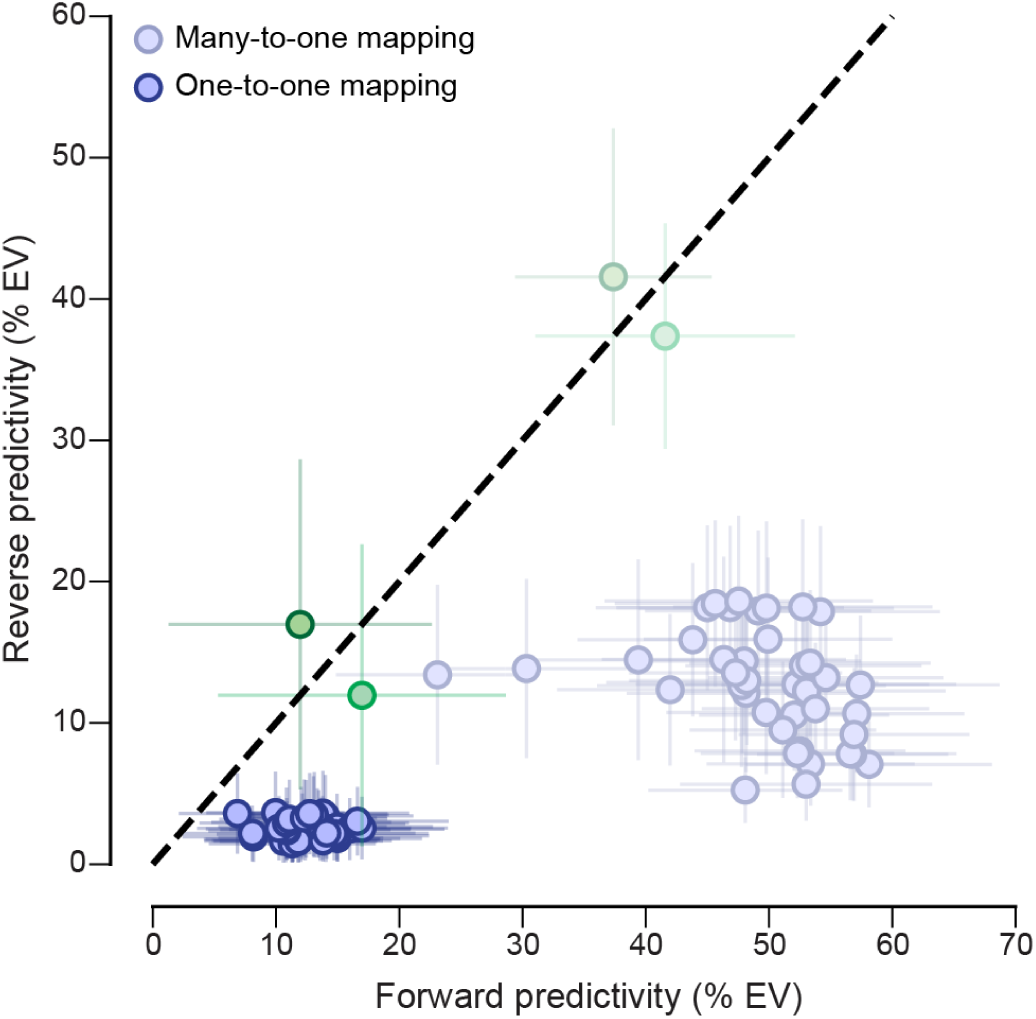
Comparison of forward vs. reverse predictivity obtained from one-to-one vs. many-to-one mapping procedures. Reverse predictivity (% explained variance) is plotted against forward predictivity for both one-to-one (dark dots) and many-to-one (light dots) mappings across models and monkeys. As expected, many-to-one mappings generally yield higher reverse EV than one-to-one mappings. However, the asymmetry remains between models and monkeys.

**Figure S3.**
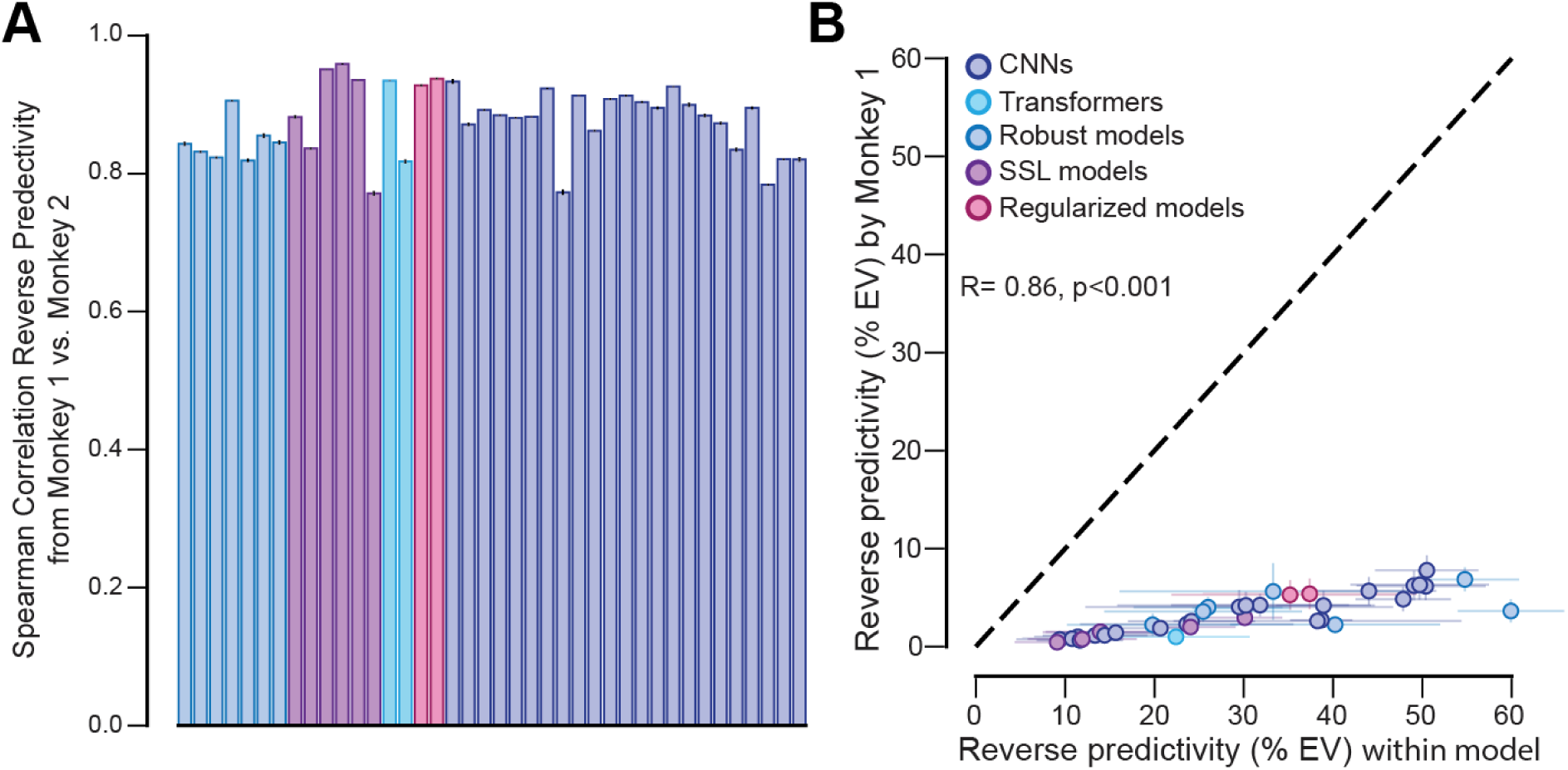
Units predictions across models. (**A**) Spearman correlations between unit-level reverse predictivity scores from two monkeys for each model. High correlations across all models indicate strong cross-subject consistency. Bars are color-coded by model type: CNNs (dark blue), Transformers (light blue), robust models (cyan), self-supervised (purple), and regularized models (red). (**B**) Scatter plot comparing reverse predictivity (% EV) when unique model units are predicted by Monkey 1 neurons (y-axis) versus predicted by units of the same model and same layer (x-axis). Each dot represents a model, colored by model class: CNNs, Transformers, Robust, SSL, and Regularized models. A strong correlation (*R = 0.86, p < 0.001*) indicates that unique units better predicted within models are also better predicted by monkeys. However, all models predict themselves better than the monkey neurons.

**Figure S4.**
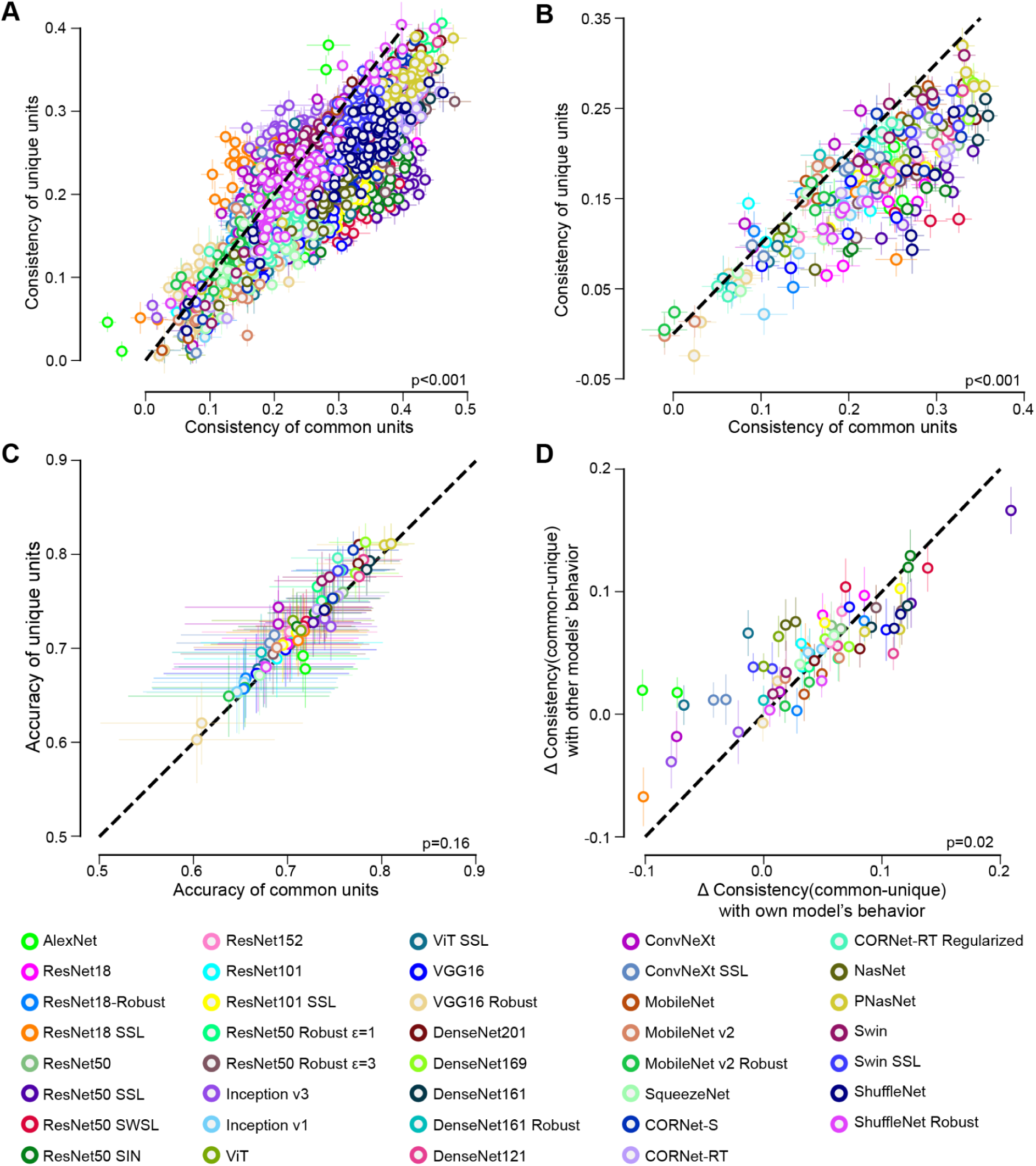
Consistency of common vs. unique units in behavioral signatures. (**A**) For each model, the consistency of image-level behavioral predictions with all other models is plotted for common vs. unique units. Each cluster of colored points corresponds to a model instance. Across models, common units exhibit higher behavioral consistency than unique ones. (**B**) Same plot as (**A**), but showing the consistency with primates’ (three monkeys and humans) image-level predictions. (**C**) Accuracy of common vs. unique units after subsampling common units to match, as closely as possible, the accuracy of the unique units. If the accuracy of the unique units falls between two subsampled values, we select the lower one for comparison. (**D**) Δ Consistency between common and unique units, computed either with respect to the behavior of the same model (within-model) or of all other models (between-model). The consistently smaller within-model deltas suggest that unique units retain some behavioral relevance specific to their source model.

**Figure S5.**
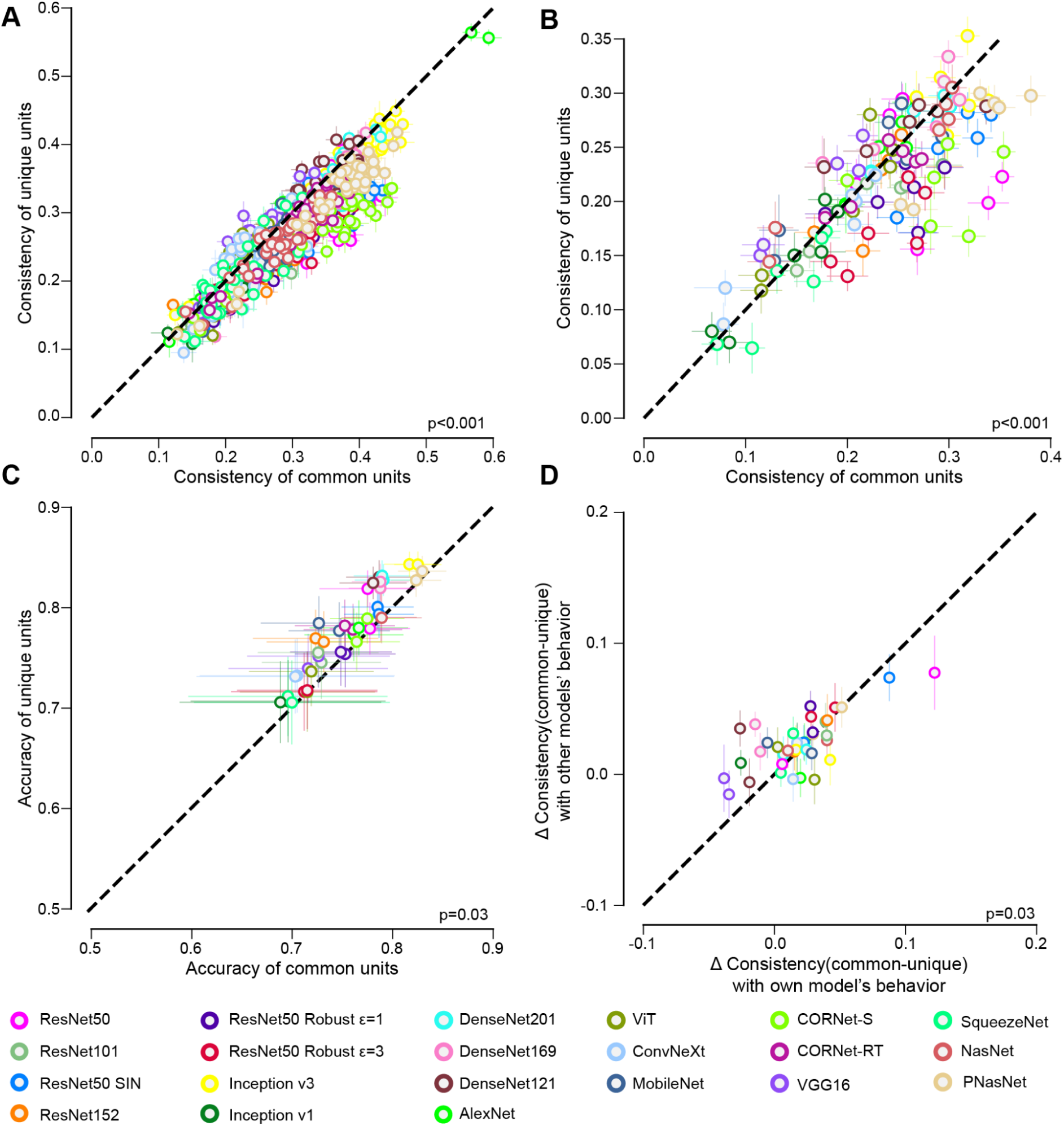
Consistency of common vs. unique units in behavioral signatures from one-to-one mapping. (**A**) For each model, the consistency of image-level behavioral predictions with all other models is plotted for common vs. unique units, selected best on reverse predictivity from one-to-one mappings. Each cluster of colored points corresponds to a model instance. Across models, common units exhibit higher behavioral consistency than unique ones. (**B**) Same plot as (**A**), but showing the consistency with primates’ (three monkeys and humans) image-level predictions. (**C**) Accuracy of common vs. unique units after subsampling common units to match, as closely as possible, the accuracy of the unique units. If the accuracy of the unique units falls between two subsampled values, we select the lower one for comparison. (**D**) Δ Consistency between common and unique units, computed either with respect to the behavior of the same model (within-model) or of all other models (between-model). The consistently smaller within-model deltas suggest that unique units retain some behavioral relevance specific to their source model.

**Table S6.**
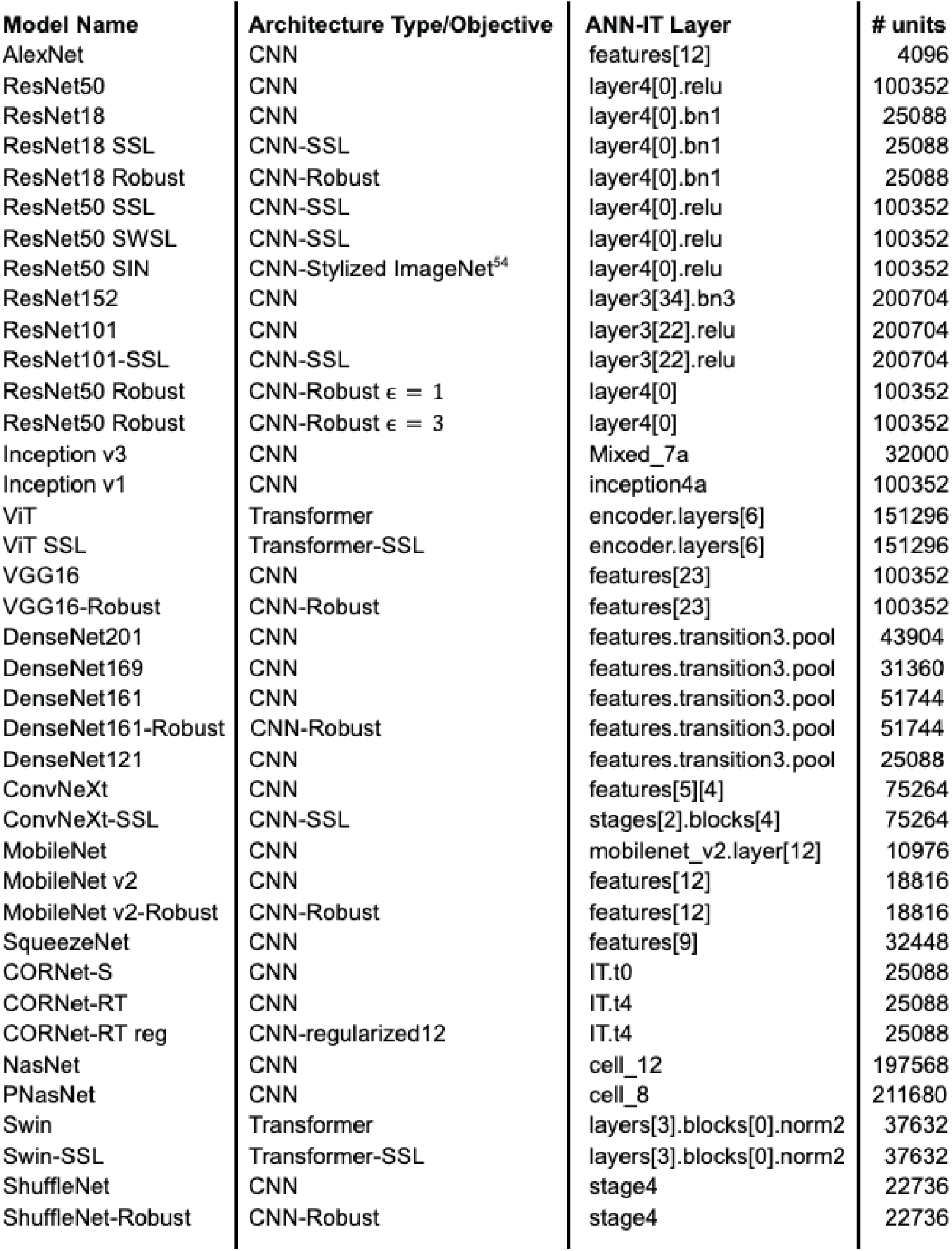
List of models with their ANN-IT layers. Exhaustive list of all the models used in this study, along with their architecture types and training objectives (if nothing is specified, Supervised Learning on ImageNet is employed), as well as their ANN-IT layer and the number of units in each of these layers.

| Model Name | Architecture Type/Objective | ANN-IT Layer | # units |
| --- | --- | --- | --- |
| AlexNet | CNN | features[12] | 4096 |
| ResNet50 | CNN | layer4[0].relu | 100352 |
| ResNet18 | CNN | layer4[0].bn1 | 25088 |
| ResNet18 SSL | CNN-SSL | layer4[0].bn1 | 25088 |
| ResNet18 Robust | CNN-Robust | layer4[0].bn1 | 25088 |
| ResNet50 SSL | CNN-SSL | layer4[0].relu | 100352 |
| ResNet50 SWSL | CNN-SSL | layer4[0].relu | 100352 |
| ResNet50 SIN | CNN-Stylized ImageNet<sup>54</sup> | layer4[0].relu | 100352 |
| ResNet152 | CNN | layer3[34].bn3 | 200704 |
| ResNet101 | CNN | layer3[22].relu | 200704 |
| ResNet101-SSL | CNN-SSL | layer3[22].relu | 200704 |
| ResNet50 Robust | CNN-Robust $\epsilon = 1$ | layer4[0] | 100352 |
| ResNet50 Robust | CNN-Robust $\epsilon = 3$ | layer4[0] | 100352 |
| Inception v3 | CNN | Mixed_7a | 32000 |
| Inception v1 | CNN | inception4a | 100352 |
| ViT | Transformer | encoder.layers[6] | 151296 |
| ViT SSL | Transformer-SSL | encoder.layers[6] | 151296 |
| VGG16 | CNN | features[23] | 100352 |
| VGG16-Robust | CNN-Robust | features[23] | 100352 |
| DenseNet201 | CNN | features.transition3.pool | 43904 |
| DenseNet169 | CNN | features.transition3.pool | 31360 |
| DenseNet161 | CNN | features.transition3.pool | 51744 |
| DenseNet161-Robust | CNN-Robust | features.transition3.pool | 51744 |
| DenseNet121 | CNN | features.transition3.pool | 25088 |
| ConvNeXt | CNN | features[5][4] | 75264 |
| ConvNeXt-SSL | CNN-SSL | stages[2].blocks[4] | 75264 |
| MobileNet | CNN | mobilenet_v2.layer[12] | 10976 |
| MobileNet v2 | CNN | features[12] | 18816 |
| MobileNet v2-Robust | CNN-Robust | features[12] | 18816 |
| SqueezeNet | CNN | features[9] | 32448 |
| CORNet-S | CNN | IT.t0 | 25088 |
| CORNet-RT | CNN | IT.t4 | 25088 |
| CORNet-RT reg | CNN-regularized12 | IT.t4 | 25088 |
| NasNet | CNN | cell_12 | 197568 |
| PNasNet | CNN | cell_8 | 211680 |
| Swin | Transformer | layers[3].blocks[0].norm2 | 37632 |
| Swin-SSL | Transformer-SSL | layers[3].blocks[0].norm2 | 37632 |
| ShuffleNet | CNN | stage4 | 22736 |
| ShuffleNet-Robust | CNN-Robust | stage4 | 22736 |

